# CeLaVi: An Interactive Cell Lineage Visualisation Tool

**DOI:** 10.1101/2020.12.14.422765

**Authors:** Irepan Salvador-Martínez, Marco Grillo, Michalis Averof, Maximilian J. Telford

**Affiliations:** Centre for Life’s Origins and Evolution, Department of Genetics Evolution and Environment, University College London, Gower Street, London, WC1E 6BT, UK; Institut de Génomique Fonctionnelle de Lyon (IGFL), École Normale Supérieure de Lyon, 32 avenue Tony Garnier, 69007 Lyon, France; Centre National de la Recherche Scientifique (CNRS), France

**Author notes:** CNAG-CRG, Centre for Genomic Regulation, Barcelona Institute of Science and Technology, Barcelona, Spain. Science for Life Laboratory, SciLifeLab, Tomtebodavägen 23, 171 65 Solna, Sweden and Department of Biochemistry and Biophysics, Svante Arrhenius väg 16C, 104 05 Stockholm, Sweden (Marco Grillo).

## Abstract

Recent innovations in genetics and imaging are providing the means to reconstruct cell lineages, either by tracking cell divisions using live microscopy, or by deducing the history of cells using molecular recorders. A cell lineage on its own, however, is simply a description of cell divisions as branching events. A major goal of current research is to integrate this description of cell relationships with information about the spatial distribution and identities of the cells those divisions produce. Visualising, interpreting and exploring these complex data in an intuitive manner requires the development of new tools. Here we present CeLaVi, a web-based visualisation tool that allows users to navigate and interact with a representation of cell lineages, whilst simultaneously visualising the spatial distribution, identities and properties of cells. CeLaVi’s principal functions include the ability to explore and manipulate the cell lineage tree; to visualise the spatial distribution of cell clones at different depths of the tree; to colour cells in the 3D viewer based on lineage relationships; to visualise various cell qualities on the 3D viewer (e.g. gene expression, cell type, tissue layer) and to annotate selected cells/clones. All these capabilities are demonstrated with four different example data sets. CeLaVi is available at http://www.celavi.pro.

## Introduction

Multicellular organisms start their development as a single cell that undergoes a coordinated process of cell division, morphogenesis and cell differentiation. The cell divisions constitute a genealogical tree —a cell lineage which can provide the framework for understanding how and when cell fate decisions are made. For many years descriptions of cell lineages have been produced by following successive cell divisions of developing organisms under the microscope (1–3). Recent years have seen development of imaging technologies that have simplified this process and made possible its application to larger numbers of cells. In parallel has been the development of genetic lineage recorders, which register the genealogical history of cells based on somatic mutations, opening up the possibility of reconstructing the entire lineage history of complex organisms (e.g. 4–8). The visualisation and integration of these complex data requires new specialised tools. Here we present CeLaVi (<u>C</u>ell <u>L</u>ine<u>a</u>ge <u>Vi</u>sualisation tool), a web based visualisation tool for exploring a cell lineage while viewing a 3D representation of the same cells, which can be annotated in any way required.

CeLaVi is an open-source web tool that requires no installation of software. Example data sets, and text and video tutorials are provided (http://www.celavi.pro/tutorial.html). The simple requirements for using CeLaVi, namely a personal computer with a modern browser, makes it easy to use by cell lineage researchers, but also opens the possibility of exploring the data emerging from the new cell lineaging technologies to the wider developmental biology community, and to the general public.

## Results and Discussion

### General description of CeLaVi

CeLaVi is built using HTML5, Ajax, jQuery, CSS and the visualisation libraries D3.js (9) and Plotly.js (10). It does not need to be installed and has been tested on the browsers and operating systems listed in Table 1.

**Table 1.** OS and browser compatibility

| OS | Version | Chrome | Firefox | Safari |
| --- | --- | --- | --- | --- |
| Linux | Ubuntu 16.04 | 87.0.4280 | 83.0 | n/a |
| MacOS | 10.15.2 | 87.0.4280 | 83.0 | 13.1.2 |
| Windows | 10 | 87.0.4280 | 83.0 | n/a |

The user interface of CeLaVi consists of two interactive spaces that are connected in real time: the ‘Lineage viewer, and the ‘3D viewer, (Figure 1). The two areas are tightly integrated so that the user can interact with the cells or cell clones on the lineage tree and observe the same cells/clones in 3D. Potential uses include exploring the cell lineage tree by collapsing/expanding individual branches; exploring the spatial distribution of the cells in 3D; visualising the spatial distribution of cell clones (descendants of an ancestral cell) by selecting an ancestral node in the lineage viewer and visualising its progeny in 3D; selecting cells in the 3D viewer to display their lineage history; visualising the clonal relationships/distances of selected cells to all the other cells in 3D.

**Fig. 1.**
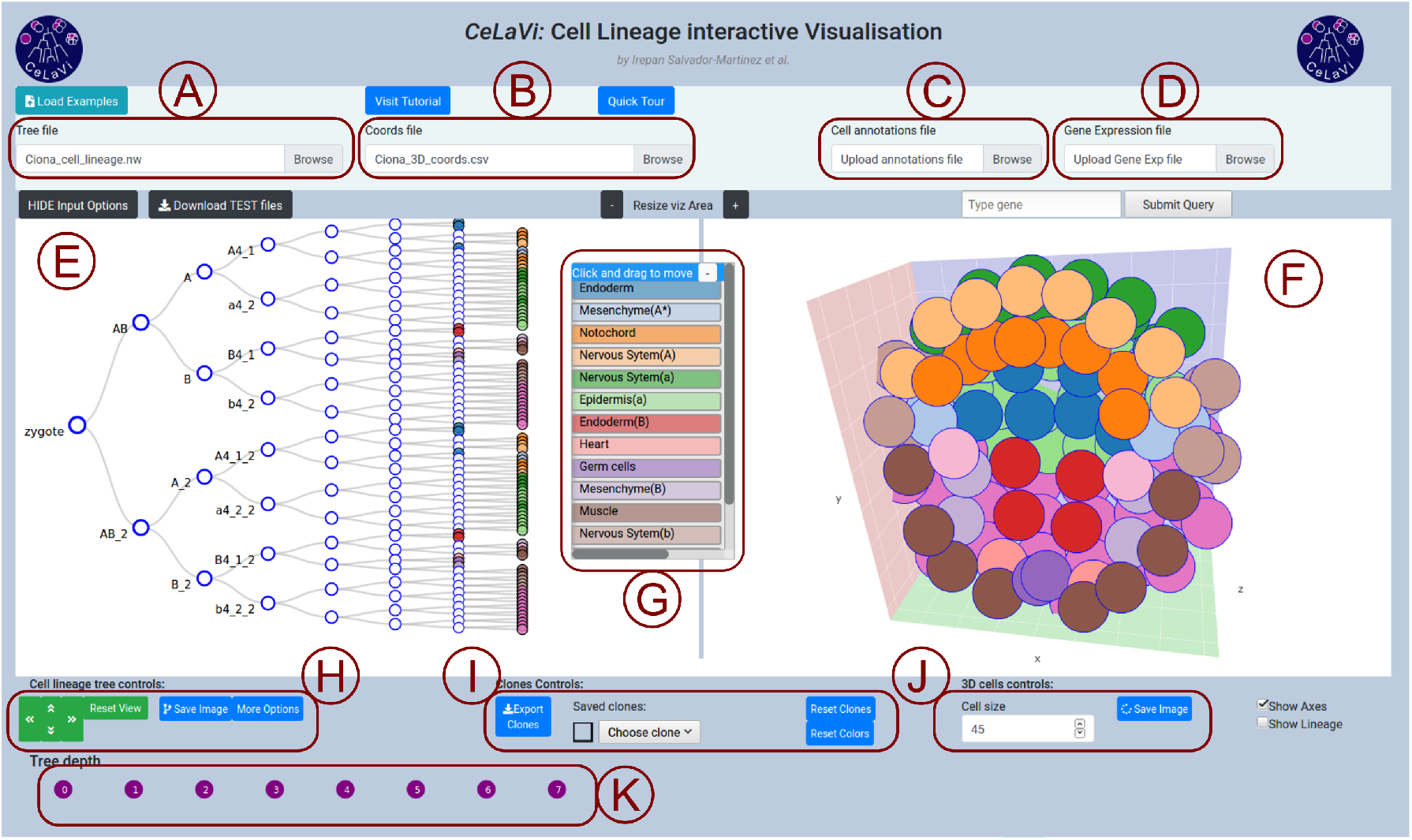
The CeLaVi user interface, loaded with the *Ciona* gastrula dataset. The main areas of the interface are labelled: A) tree file input; B) cell coordinates file input; C) cell annotations file input; D) gene expression file input; E) lineage viewer; F) 3D viewer; G) cell annotations table; H) cell lineage tree controls; I) clones controls; J) 3D cells controls; K) tree depth area. Cells show cell fates with a colour code based on the cell annotations table. For more details visit the tutorial (http://www.celavi.pro/tutorial.html).

### Input/Output

To visualise a cell lineage tree and to relate it to a 3D representation of the positions of each cell, the user must provide two files: one containing the lineage tree and one containing the 3D coordinates of the same cells (with identical labels).

The lineage tree file describes the topology of the tree either in Newick or Json format. When uploading a file (Figure 1A), CeLaVi automatically identifies the format of the lineage tree file and displays the tree in the Lineage tree viewer (Figure 1E). To ensure the format of the lineage tree is correct, Json and Newick files are parsed with the Javascript standard method JSON.parse() or with the newick-tools software (11; available at https://github.com/xflouris/newick-tools), respectively. Branch lengths can be recorded in the input file, either as ‘absolute, branch lengths (distances of each node from the root) or as ‘relative, blanch lengths (distances between nodes of the tree).

To visualise the same cells in the 3D viewer, the user needs to upload a file of the 3D coordinates of cells (Figure 1B) in comma-separated-value (csv) format. The first row of the file is a header specifying the identities of each column (“cell”, “X”, “Y”, and “Z”) and each subsequent row giving the name/ID and 3D coordinates (X, Y, Z) of each cell. The cells are represented in an interactive visualisation (Figure 1F) in the 3D viewer. The image of the lineage tree or the 3D cells, as currently displayed in the visualisation areas, can be exported as an image file (in SVG or PNG format) at any time.

For MaMuT (12) users, it is possible to convert a MaMuT single XML output file into CeLaVi lineage and 3D coordinates input files (in json and csv formats respectively) using the python script “mamut_to_celavi” by Ko Sugawara, available at https://github.com/ksugar/celavi-tools.

### Lineage tree viewer

CeLaVi’s lineage tree viewer (Figure 1E) has been built to be scalable in order to accommodate very large trees while retaining tree legibility. To reduce the computational burden, CeLaVi has an option to plot only a subset of the tree branches. This option is automatically activated when the number of cells in the tree is greater than 500. This function is only applicable to lineage trees that are not completely resolved, i.e. with internal nodes giving rise to more than two branches (polytomies). The CeLaVi lineage viewer also allows the user to focus on specific branches of the lineage tree by expanding, collapsing or pruning selected branches.

Interacting with each cell in the lineage tree (including ancestral cells at nodes within the tree) allows the user to display a number of functions: hovering the pointer over a cell will display the cell ID, total number of descendants and number of descendants on the 3D viewer, and highlight the corresponding cell (or clone) in the 3D viewer; clicking cells marks the corresponding cells/clones with distinct colours in the 3D viewer; right-clicking allows a choice of expanding, collapsing, pruning or saving the selected cells/clones. For cell lineages containing branch length information (reflecting the relative timing of cell divisions), it is possible to change the visualisation mode from a cladogram with fixed branch lengths, to a branch-length based representation.

### 3D viewer

CeLaVi’s 3D viewer renders cells as spheres. The user can zoom in/out using the scroll wheel of the mouse, rotate the image by left-clicking and dragging, panning the 3D scatterplot by right-clicking and dragging. Clicking any cell in the 3D viewer highlights its full lineage, in the lineage viewer. The “Show lineage” option allows the user to visu-alise the degree of lineage relationship of all cells relative to one selected cell showing the nested set of clones to which the cell belongs.

### Visualising cell clones

The spatial distribution of cell clones in CeLaVi can be visualised by clicking on any ancestral cell in the lineage tree. This marks with a random colour all the descendants of that cell on the 3D viewer. Multiple clones can be thus selected and visualised at the same time with unique colours.

All the clones originating at a given tree depth (i.e. after a certain number of divisions or, for trees with branch lengths, existing at a certain time point) can be marked with distinct colours by using the “Tree depth” area below the lineage tree (Figure 1K).

CeLaVi allows the user to save selected cells or cell clones, with customized annotations, in a dropdown list that can be used later to visualise them (Figure 1I). Sets of cells/clones can also be saved as a .csv file that can later be reloaded as a cell annotation file (see below).

### Visualising cell annotations and gene expression

Two additional files can be loaded to provide information associated with the cells in the two viewer windows. Cell type information (or indeed any other cell annotation) can be uploaded as a csv file using the “Cell annotations file” input box (Figure 1C). The information from this file is displayed as an interactive table with the annotations as coloured rows (Figure 1G). For example, selecting the “muscle” annotation in the Ciona dataset, labels the muscle cells in both the 3D viewer and the lineage viewer.

Gene expression data (Figure 1D), can be uploaded as a .csv file using the “Gene expression file” input box. To see the relative expression levels of a gene of interest, the user types the gene name in the search box (which has an autofill function) and clicks “submit”. The gene expression levels will be represented on both the 3D and lineage viewers using a heatmap. Figure 2 shows the expression levels of gene Sox14 in the C. *intestinalis* gastrula.

**Fig. 2.**
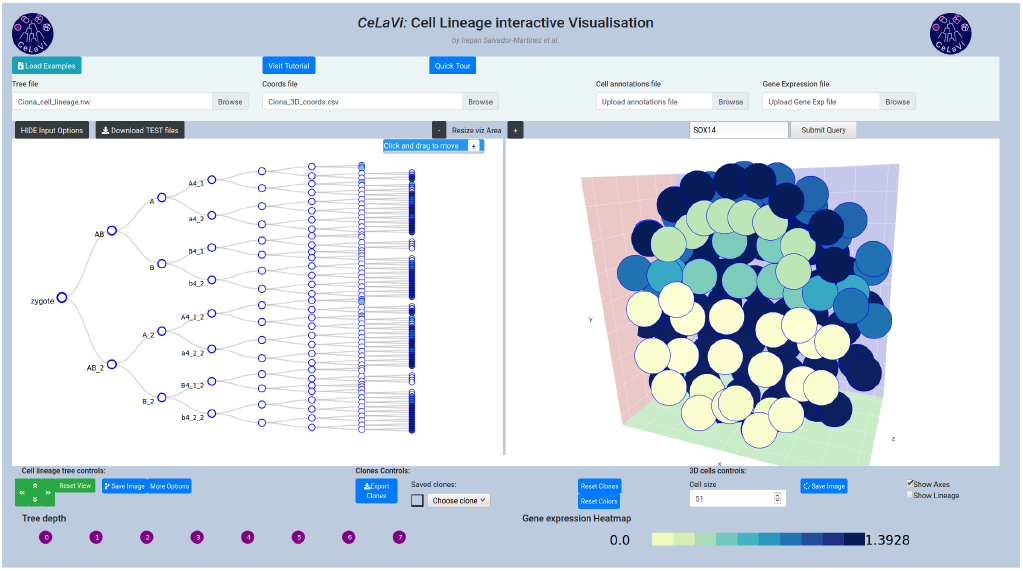
*Ciona* gastrula example dataset showing the expression pattern of gene Sox14 as a heatmap (colour code below the 3D viewer area).

### Example datasets

We include 4 example datasets that demonstrate CeLaVi’s features (see Table 2).

**Table 2.** Example datasets available in CeLaVi.

|  | Cell lineage (format) | 3D coords | Branch lengths | Cell annotations | Gene expression |
| --- | --- | --- | --- | --- | --- |
| *C. elegans* L1 larva | yes (newick) | yes | yes (relative) | yes | no |
| *Ciona* gastrula | yes (newick) | yes | no | yes | yes |
| *Parhyale* limb | yes (json and newick) | yes | yes (absolute) | no | no |
| *In silico* Organoid | yes(json) | yes | no | no | no |

### Caenorhabditis elegans L1 larva

The embryonic cell lineage of the nematode *Caenorhabditis elegans* was first described by Sulston and collaborators in 1983 (2). The data used for the cell lineage and the cell types were extracted from a json file made available by Nikhil Bhatla at wormweb.org/celllineage. The complete cell lineage was trimmed back to the L1 larval stage. The 3D coordinates come from the study of Long and collaborators (13), where they mapped the positions of 357 cells (out of the 558 cells) of the L1 larval stage, using confocal image stacks from 15 individual worms.

### Ciona intestinalis gastrula

The cell lineage of the ascidian *Ciona intestinalis* was described by Conklin in 1905 (14). The cell lineage example/test file (in newick format) was created using Conklin’s lineage and nomenclature, from the zygote to the 110-cell stage. The 3D coordinates of the cells in the Ciona gastrula were obtained from a reconstructed 3D embryo model (Mid_112-cell_stage_Amira_1.obj file; 15), available in the Aniseed database (16). The 3D embryo model contains information in cell shapes. We used the cell centroid to represent each cell as a single point in 3D space. The gene expression data were obtained from the single-cell sequencing dataset published by Levine and collaborators (17). We used the raw expression data matrix (ex-pression_matrix_C110.1.tsv), containing the expression levels of 15,307 genes in 1,731 cells, together with the associated metadata file (C110.1.clusters.upload.rename.1.txt) that contains the cell type identity of each cell at the early gastrula stage. We used the R software version 3.6.3 (18) with the Seurat package version 3.1.4 (19) to identify the 500 genes with the most variable expression between cells and to obtain their average expression values for each cell. The gene names correspond to human gene IDs, derived from the *Ciona intestinalis* genome assembly KH2012 with NCBI Gene Model to Best Blast Hit mapping, available in the Aniseed database. Only the gene ID with the highest Blast e-value is shown.

### Parhyale hawaiensis limb

The cell lineage and 3D coordinates of the limb of the crustacean *Parhyale hawaiensis* were obtained by Pavlopoulos and collaborators (12). Transgenic fluorescently-labeled embryos were imaged with multi-view light-sheet microscopy at high spatiotemporal resolution over several days of embryogenesis. The cell lineage was recon-structed with the aid of the MaMuT software.

### In silico Organoid

The “organoid” dataset comes from a simulation of morphogenesis using the yalla software (20). The organoid is the product of a simulation of branching morphogenesis with epithelium and mesenchyme (only the epithelial cells are recorded in this example). The simulation was kindly provided by Miquel Marín-Riera, coauthor of the yalla software and is based on the example “branching.cu” available in https://github.com/germannp/yalla.

### Comparison with other software

CeLaVi is, to our knowledge, the first interactive web-based visualisation of cell lineages. It was inspired by web based visualisations of phylogenetic trees, such as phylo.io and Evolview3 (21, 22). The most similar existing implementation of some of these features is found in Morphonet (23) which is a web-server software for visualising morphological data. Morphonet shares some commonalities with CeLaVi, but cell lineage viewing is not its main focus, and the interactive manipulation of the cell lineage data as well as its integration with the spatial information is limited. CeLaVi’s simpler representation of cells in 3D enables very fast rendering of visualisations and this is essential to the exploration of the data.

## Data Availability

CeLaVi is freely available at: http://www.celavi.pro. It is an open source software, with the source code available at https://github.com/irepansalvador/CeLaVi.

The example datasets described here can be downloaded directly from the web server as a compressed file (zip format). A complete tutorial can be found at http://www.celavi.pro/tutorial.html.

## Acknowledgements

We thank Anastasios Pavlopoulos for kindly providing lineage data and cell coordinates for the *Parhyale* limb dataset, Chen Cao and Michael Levine for kindly providing the scRNAseq data for the *Ciona* gastrula, Miquel Marín-Riera for kindly providing the organoid simulation, and Ko Sugawara for producing the script to input MaMuT data. We thank Tomáš Flouri for fruitful early discussions on the implementation of CeLaVi and general guidance. We also want to thank the members of the Telford, Oliveri and Averof labs for testing CeLaVi and providing useful feedback.

